# Identification of tumor-intrinsic drivers of immune exclusion in acral melanoma

**DOI:** 10.1101/2023.08.24.554717

**Authors:** Ryan C. Augustin, Sarah Newman, Aofei Li, Marion Joy, Maureen Lyons, Mary Pham, Peter C. Lucas, Kate Smith, Cindy Sander, Brian Isett, Diwakar Davar, Yana G. Najjar, Hassane M. Zarour, John M. Kirkwood, Jason J. Luke, Riyue Bao

## Abstract

**Background:** Acral melanoma (AM) has distinct characteristics as compared to cutaneous melanoma and exhibits poor response to immune checkpoint inhibitors (ICI). Tumor-intrinsic mechanisms of immune exclusion have been identified in many cancers but less studied in AM.

**Methods:** We characterized clinically annotated tumors from patients diagnosed with AM at our institution in correlation with ICI response using whole transcriptome RNAseq, whole exome sequencing, CD8 immunohistochemistry, and multispectral immunofluorescence imaging. A defined interferon-γ-associated T cell-inflamed gene signature was used to categorize tumors into non-T cell-inflamed and T cell-inflamed phenotypes. In combination with AM tumors from two published studies, we systematically assessed the immune landscape of AM and detected differential gene expression and pathway activation in a non-T cell-inflamed tumor microenvironment (TME). Two single-cell(sc) RNAseq AM cohorts and 11 bulk RNAseq cohorts of various tumor types were used for independent validation on pathways associated with lack of ICI response. In total, 892 specimens were included in this study.

**Results:** 72.5% of AM tumors showed low expression of the T cell-inflamed gene signature, with 23.9% of total tumors categorized as the non-T cell-inflamed phenotype. Patients of low CD3^+^CD8^+^PD1^+^ intratumoral T cell density showed poor prognosis. We identified 11 oncogenic pathways significantly upregulated in non-T cell-inflamed relative to T cell-inflamed TME shared across all three acral cohorts (MYC, HGF, MITF, VEGF, EGFR, SP1, ERBB2, TFEB, SREBF1, SOX2, and CCND1). scRNAseq analysis revealed that tumor cell-expressing pathway scores were significantly higher in low vs high T cell-infiltrated AM tumors. We further demonstrated that the 11 pathways were enriched in ICI non-responders compared to responders across cancers, including acral melanoma, cutaneous melanoma, triple-negative breast cancer, and non-small cell lung cancer. Pathway activation was associated with low expression of interferon stimulated genes, suggesting suppression of antigen presentation. Across the 11 pathways, fatty acid synthase and CXCL8 were unifying downstream target molecules suggesting potential nodes for therapeutic intervention.

**Conclusions:** A unique set of pathways is associated with immune exclusion and ICI resistance in AM. These data may inform immunotherapy combinations for immediate clinical translation.

## Background

Acral melanoma (AM) is a unique subtype portending poor prognosis as compared to cutaneous melanoma (CM).^1^ Acral tumors arise in sun-shielded locations (e.g. palms/soles), often leading to delayed medical attention and advanced stage at diagnosis. While UV radiation-induced CM demonstrates high tumor mutational burden (TMB), neoantigenicity, and frequency of *BRAF* mutations, AM has lower TMB and rarely demonstrates *BRAF* mutations.^2^ AM is also characterized by lower CD8^+^ T cell infiltration and Programmed Death Ligand-1 (PD-L1) expression, hallmarks of a non-T cell-inflamed tumor microenvironment (TME). Whereas CM is associated with an approximately 40% objective response rate (ORR) in the treatment naïve metastatic setting to anti-PD1 monotherapy, reports of response in AM are quoted as less than 20%.^3^ A single-cell expression analysis of AM tumors demonstrated the expression of certain inhibitory checkpoints but primarily highlighted the relative lack of immune cell infiltrate.^4^

An interferon-γ (IFNg) activated TME is strongly associated with response to immune checkpoint inhibitors (ICI),^5^ yet most clinical biomarkers (PD-L1, TMB, tumor infiltrating lymphocyte (TIL) levels) fail to completely encompass this inflammatory phenotype. A more comprehensive T cell-inflamed gene signature has been shown to strongly correlate with IFNg signaling and ICI response.^6^ Additionally, this signature provides a model whereby ICI resistance mechanisms can be identified by comparing non-T cell-inflamed versus T cell-inflamed tumors.^7^ In CM, the Wnt/β-catenin pathway was identified as a driver of immune exclusion in non-T cell-inflamed tumors.^8^ This approach has further nominated other pathways (e.g., MYC activation and PTEN loss) associated with suppression of antigen presentation and cytolytic activity across human cancers.^7^

To inform combination immunotherapy drug development in AM, we sought to identify tumor-intrinsic drivers of immune exclusion. To this end, we collected a set of acral tumors from individuals exposed to ICI and investigated molecular mediators of non-T cell-inflamed AM. Through this approach, we have nominated a series of molecular targets as potential combination strategies to enhance immune checkpoint blockade in AM.

## Methods

A full description is supplied in the **Supplementary Methods**. In brief: Formalin-fixed, paraffin-embedded (FFPE) baseline tumor tissues along with demographic, histopathological, and treatment data were collected from the melanoma biospecimen bank of UPMC Hillman Cancer Center (UPMC) (20 patients, 14 with ICI response evaluable; **Table S1**), according to an institutional review board-approved protocol (No. 20090109). Bulk RNAseq, whole exome sequencing (WES), immunohistochemistry (IHC), and multispectral immunofluorescence (mIF) imaging with PhenoImager^TM^ HT (Akoya Biosciences) was performed for each sample. Baseline AM tumors from two additional published studies (NW and MIA cohorts), two scRNAseq AM cohorts (Li *et al*. and Zhang *et al*. cohorts), and 11 tumor cohorts from ICI-treated patients with bulk RNAseq and clinical data available were also analyzed (**Table S2**). In total, 892 specimens were analyzed across all cohorts. Acral samples were clustered into T cell-inflamed cohorts using a defined T cell-inflamed gene signature;^6^ these clusters were used for differential gene expression detection by limma voom with precision weights and pathway activation prediction by Ingenuity causal network. For scRNAseq cohorts, high and low T cell-infiltrated tumors were defined by fraction of T cells over all cells profiled in TME. Statistical analysis was performed using R (v4.1.2) and Bioconductor (release 3.14), with significance level at 0.10. Benjamini-Hochberg (BH) Procedure was used to adjust multiple comparisons.

## Results

### Characterization of the acral melanoma immune landscape

We assembled baseline tumor RNAseq data from 109 patients in three independent AM cohorts, including UPMC (*n*=20), Northwestern University (NW; *n*=22) (Shi *et al*., **Table S2**), and Melanoma Institute of Australia (MIA; *n*=67) (Newell *et al*., **Table S2**). We generated RNAseq and WES data as well as clinical annotation for specimens from 20 patients in the UPMC cohort and validated our findings in 89 patients from two publicly available AM datasets (NW and MIA). Demographic, histopathologic, treatment, and ICI response variables for the UPMC cohort are listed in **Table S1**.

To investigate the immune landscape of AM, we used the T cell-inflamed gene expression signature^7^ to categorize tumors into T cell-inflamed and non-T cell-inflamed groups, with the rest as intermediate. The overall analysis workflow is provided in **Fig. S1**. Since each dataset was generated at a different institution with variations in experimental approaches, strong technical batch effect exists which cannot be fully corrected by computational methods. Therefore, we took an ensemble approach to analyze each dataset independently and then cross-reference results to identify consistent patterns relevant to the biological phenotypes. We detected high expression of T cell signature genes (T cell-inflamed) in 20%, 22.7%, and 31.3% tumors from UPMC, NW, and MIA cohorts, respectively, consistent with T cell infiltration data from AM single-cell cohorts.^4,9^ Non-T cell-inflamed tumors were identified in 20% (UPMC), 22.7% (NW), and 25.4% (MIA) samples (**Fig. 1A**). The T cell-inflamed tumors showed upregulated *CD8A* gene expression as well as a higher TMB (UPMC, 121.7±27.8; MIA, 121.6±37.0) compared to the non-T cell-inflamed tumors (UPMC, 51.5±6.5; MIA, 100.4±17.3) (**Fig. 1A**). The NW cohort does not have TMB data available. Digital immune deconvolution of RNAseq data revealed significantly elevated CD8^+^ T cell populations in T cell-inflamed versus non-T cell-inflamed TME in all three cohorts (*P*<0.05) (**Fig. 1B**). We detected no significant correlation between the T cell-inflamed gene expression and typical driver mutations of melanoma (*BRAF, NRAS, NF1*) (**Fig. S2A**).

**Figure 1.**
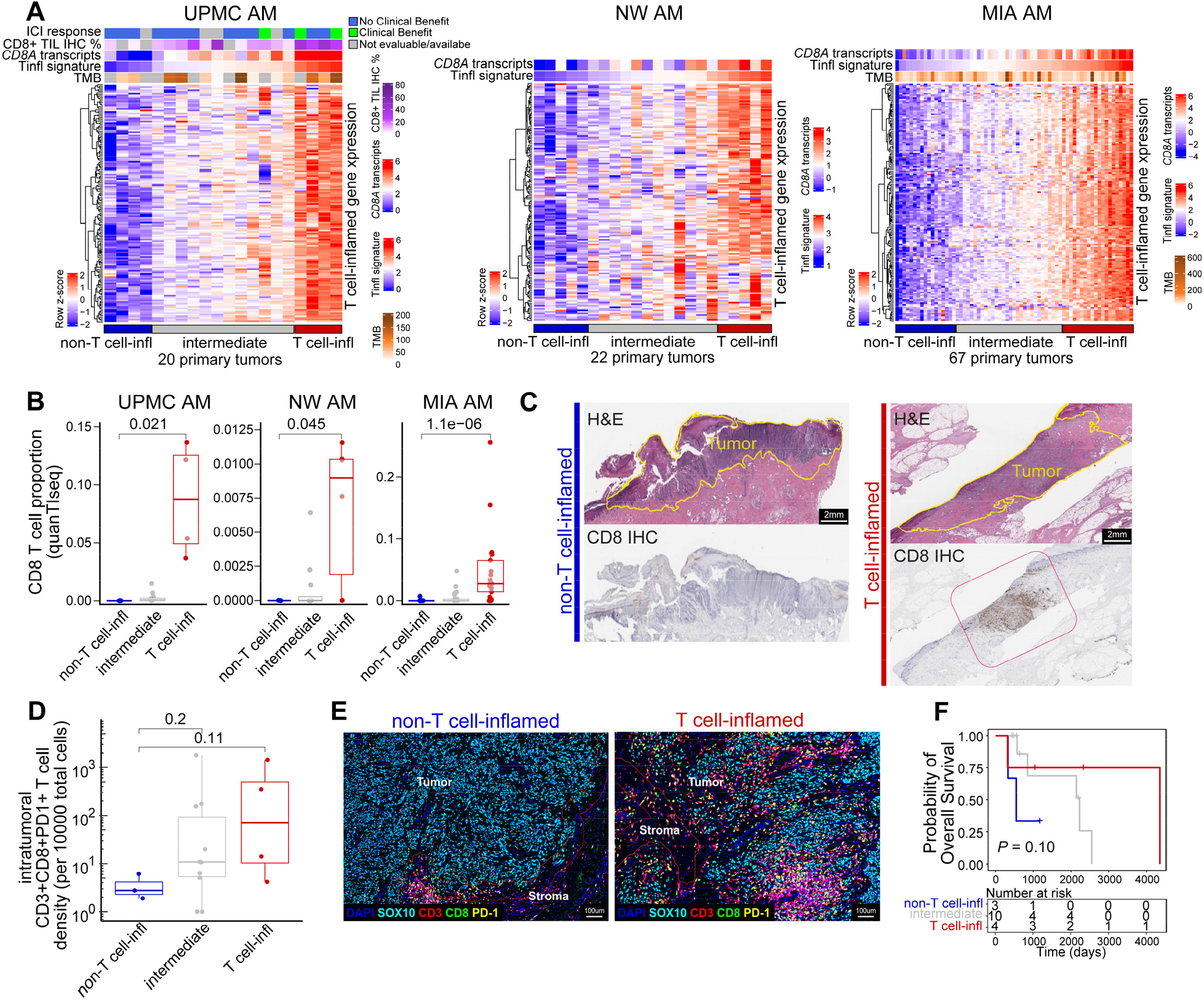
Immune landscape of acral tumors across three cohorts (UPMC, NW, MIA). (**A-B**) *n*=109 samples shown with RNAseq data from all AM cohorts. Hierarchical clustering based on the expression of a defined T cell-inflamed gene signature identifies T cell-inflamed, intermediate, and non-T cell-inflamed tumor groups. Horizontal annotation bars above the heatmaps represent ICI response, CD8^+^ tumor lymphocyte infiltrates by IHC, CD8A gene expression and T cell-inflamed gene expression by RNAseq, and total TMB by WES, when data is available. (**B**) CD8^+^ T cell fraction in the three tumor groups via digital immune deconvolution from RNAseq data. (**C-G**) *n*=17 shown with tumor specimens available for imaging experiments from the UPMC cohort. (**C**) CD8^+^ IHC staining in representative images of a non-T cell-inflamed and T cell-inflamed tumor. Pink circle highlights heavy CD8 IHC staining. (**D**) Intratumoral CD3^+^CD8^+^PD1^+^ T cell infiltrates in the three tumor groups via multispectral immunofluorescence imaging. (**E**) Representative immunofluorescence images of non-T cell-inflamed and T cell-inflamed acral tumors. (**F**) Kaplan-Meier survival curves stratified by tumor groups, with p-value shown for testing T cell-inflamed and intermediate tumor groups versus non-T cell-inflamed tumor group. P-values were computed by two-sided Wilcoxon test in **B** and **D**, and by log-rank test in **F**.

In support of the *in silico* findings, we performed IHC on 17 tumor specimens available from the UPMC cohort, demonstrating that higher CD8^+^ tumor infiltrating lymphocyte (TIL) percentage is significantly correlated with a T cell-inflamed phenotype (Spearman’s correlation ρ=0.51, *P*=0.039) (**Fig. S2B**; representative images shown in **Fig. 1C**). A clear trend was also demonstrated by mIF imaging for increased intratumoral CD3^+^CD8^+^PD1^+^ T cell infiltrates in the T cell-inflamed versus non-T cell-inflamed (*P*=0.11) or intermediate (*P*=0.20) UPMC samples (**Fig. 1D**), further validating the immunophenotypic groups used for downstream analysis. Representative non-T cell-inflamed and T cell-inflamed mIF images are shown in **Fig. 1E**. Overall, mIF data indicate these tumors are primarily “immune excluded” with a significantly higher proportion of CD3^+^CD8^+^PD1^+^ T cells residing in the tumor edge (**Fig. S2C, S2D**). Furthermore, whether measured in the intratumoral, tumor edge, or stromal space, a higher proportion of CD3^+^CD8^+^PD1^+^ T cells correlated with improved overall survival (OS) (**Fig. S2E**).

We and other groups have previously demonstrated that the T cell-inflamed gene signature predicts response to ICI across human solid tumors.^5^ Within the UPMC AM dataset, we observed 100% of tumors from patients receiving clinical benefit fell in the upper 50^th^ percentile of T cell-inflamed gene expression. Seventeen patients had survival data available and were included for assessing the association of the T cell-inflamed gene expression profile with outcome. Patients harboring T cell-inflamed or intermediate tumors also experienced improved OS versus patients with non-T cell-inflamed tumors (*P*=0.10, by log-rank test; **Fig. 1F**). T cell-inflamed gene expression as a continuous variable was significantly associated with longer OS (*P*=0.048, hazard ratio (HR)=0.49, by Cox PH univariate model), and remained significant after adjustment for age, gender, or TMB (*P*=0.041, HR=0.32, by Cox PH multivariable model; **Fig. S2F**).

### Tumor-intrinsic oncogenic transcriptional programs are associated with immune exclusion across three acral melanoma datasets

To investigate the transcriptional programs that potentially drive the non-T cell-inflamed phenotype, we analyzed tumors from the UPMC, NW, and MIA cohorts. We took an unbiased approach^7,8^ by comparing the whole transcriptome expression of non-T cell-inflamed versus T cell-inflamed tumors using bulk RNAseq. Considering batch effect, we first performed the comparison within individual cohorts independently and then reported pathways that were common to all three cohorts. There were 2363, 1555, and 2965 differentially expressed genes (DEGs) between the two tumor groups in UPMC, NW, and MIA cohorts, respectively (*P*<0.05, fold change ≥ 1.5 or ≤ -1.5).

While common DEGs exist among all three cohorts (**Tables S3**), we sought to identify pathways that may reflect the actual biologic phenotype relevant for translational targeting rather than individual genes. To gain insights into the functional mechanisms collectively driven by DEGs, we predicted the activation of upstream regulators (pathways) based on the aggregate change of expression from downstream target molecules (encoded by DEGs), using causal network analysis with Ingenuity Knowledge Base^10^ (Qiagen Inc.) (**Tables S4-S6**). We identified 393, 149, and 70 pathways predicted to be activated in non-T cell-inflamed versus T cell-inflamed tumors from the UPMC, NW, and MIA cohorts, respectively (z-score ≥ 0.9, *P*<0.05) (**Fig. 2A**). Among those, we found known drivers of immune exclusion such as TGFB, HIF1A, and CTNNB1 (β-catenin).^8,11,12^ Upon cross-referencing results across all three cohorts, we identified 11 activated pathways consistently detected in the non-T cell-inflamed TME, including MYC, HGF, MITF, VEGF, EGFR, SP1, ERBB2, TFEB, SREBF1, SOX2, and CCND1 (**Fig. 2A**). Including the intermediate T cell-inflamed tumors in an additional investigation, we demonstrated that the pathway expression scores of all 11 pathways were significantly and inversely correlated with the T cell-inflamed signature in a continuous manner (**Fig. 2B**).

**Figure 2.**
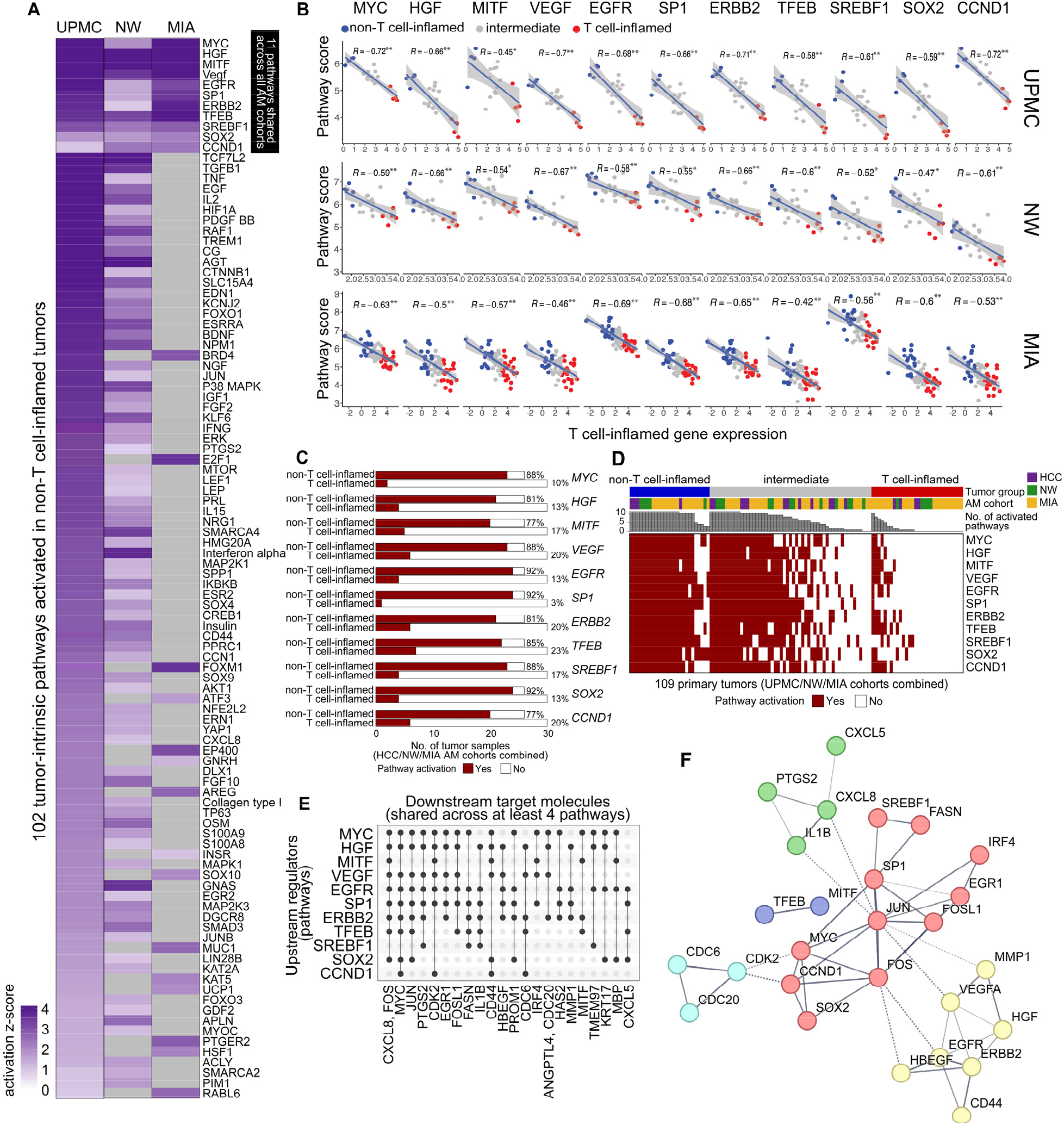
Activation of 11 oncogenic pathways in non-T cell-inflamed versus T cell-inflamed acral tumors across three cohorts (UPMC, NW, MIA). (**A**) Pathways activated in non-T cell-inflamed versus inflamed UPMC tumors and one or both of the NW and MIA cohorts. 102 pathways with a z-score of at least 0.9 at *P*<0.05 in at least one cohort are shown. Eleven shared pathways are identified across the three cohorts (shown on top, solid side bar). Each row represents one pathway. (top to bottom) pathways were sorted by z-score high to low in the UPMC cohort. (**B**) Correlation between pathway expression scores and the T cell-inflamed gene signature. All samples are shown; *n*=20 in UPMC, *n*=22 in NW, and *n*=67 in MIA. Each data point represents one tumor; color denotes the T cell-inflamed (red), non-T cell-inflamed (blue), and intermediate groups (grey). Spearman’s correlation coefficient ρ and FDR-adjusted p-values are shown for each pathway. Linear regression was shown with 95% confidence bands. (**C**) The proportion of tumors harboring activation of each pathway in non-T cell-inflamed and T cell-inflamed groups. x-axis shows the number of tumors with all three acral cohorts combined. (**D**) Correlation between the T cell-inflamed gene signature and activation of each pathway at a continuous scale. (left to right) tumor samples were sorted by by higher to lower number of activated pathways within each group. *n*=109 tumors are shown. (**E**) Downstream target molecules shared among the 11 pathways. Molecules shared by at least four pathways are shown. (**F**) Protein-protein functional network of the 11 pathways and downstream target molecules from **E**. Nodes with at least one connection are shown, from STRING functional protein association networks (confidence score >0.4; active interaction sources as “Experiments”, “Databases”, and “Co-expression”). Line thickness indicates the strength of data support. Nodes were clustered by graph-based Markov Cluster Algorithm (MCL) (inflation parameter = 2.5), with color indicating each cluster and dotted lines indicating edges between clusters. P-values were computed by Spearman’s correlation in **B**, with denotation: * P <0.05; ** P <0.01; *** P <0.001 after FDR adjustment for multiple comparisons.

From a precision immuno-oncology perspective, we sought to investigate the spectrum of pathway activation in individual patient’s tumors, and calculated an activation score for each pathway using previously described methods.^7^ Our result demonstrated that the 11 pathways operate in a relatively homogenous manner, with activation of each pathway detected in at least 77% of non-T cell-inflamed tumors, compared to 23% or less of the T cell-inflamed tumors (**Fig. 2C**). Given the high degree of pathway co-activation in individual tumors (**Fig. 2D**), we analyzed for shared downstream mechanisms across the 11 gene pathways. Both CXCL8 (also known as interleukin-8; IL-8) and FASN (fatty acid synthase) were observed to be among the top target molecules shared across pathways (**Fig. 2E**), altogether forming a functional network of associated proteins regulating cell cycle, fatty acid biosynthesis, and cytokine production (**Fig. 2F**).

### Single cell analysis of two acral melanoma cohorts confirms tumor cell-intrinsic expression of immunosuppressive pathways with significant upregulation in low T cell-infiltrated tumors

To assess the cellular derivation of the 11 shared pathways, we analyzed a cohort of AM scRNAseq tumors (**Fig. 3A**) (Li *et al*., **Table S2**). Eight of 11 pathways (MYC, MITF, EGFR, SP1, ERBB2, SREBF1, SOX2, and CCND1) were included for downstream analysis after removing redundant target molecules and requiring at least five unique molecules per pathway. Tumor cells were observed as the predominant source of expression for all pathways (**Fig. 3B**). The expression of each pathway was significantly elevated in tumor cells of low T cell-infiltrated relative to those of high T cell-infiltrated samples, including a combined score of all pathways (**Fig. 3C**). Additionally, tumor cells of low T cell-infiltrated samples showed a significant down-regulation in the expression of 108 DEGs enriched in IFN stimulating functions, consistent with suppression of type I and II IFN signaling (**Fig. 3D-3E; Table S7**). An additional validation cohort of five scRNAseq AM samples (Zhang *et al*., **Table S2**) similarly showed a predominance of pathway expression from tumor cells. More specifically, significant upregulation of MITF, SP1, ERBB2, SOX2, and CCND1 pathways in low T cell-infiltrated relative to high T cell-infiltrated samples (the latter exhibiting tumor clusters enriched for type I and II IFN signaling) was observed (**Fig. S3A-S3D**). Altogether, these data highlight a tumor-intrinsic source of these immunosuppressive pathways leading to a TME state associated with suppressed antigen presentation.

**Figure 3.**
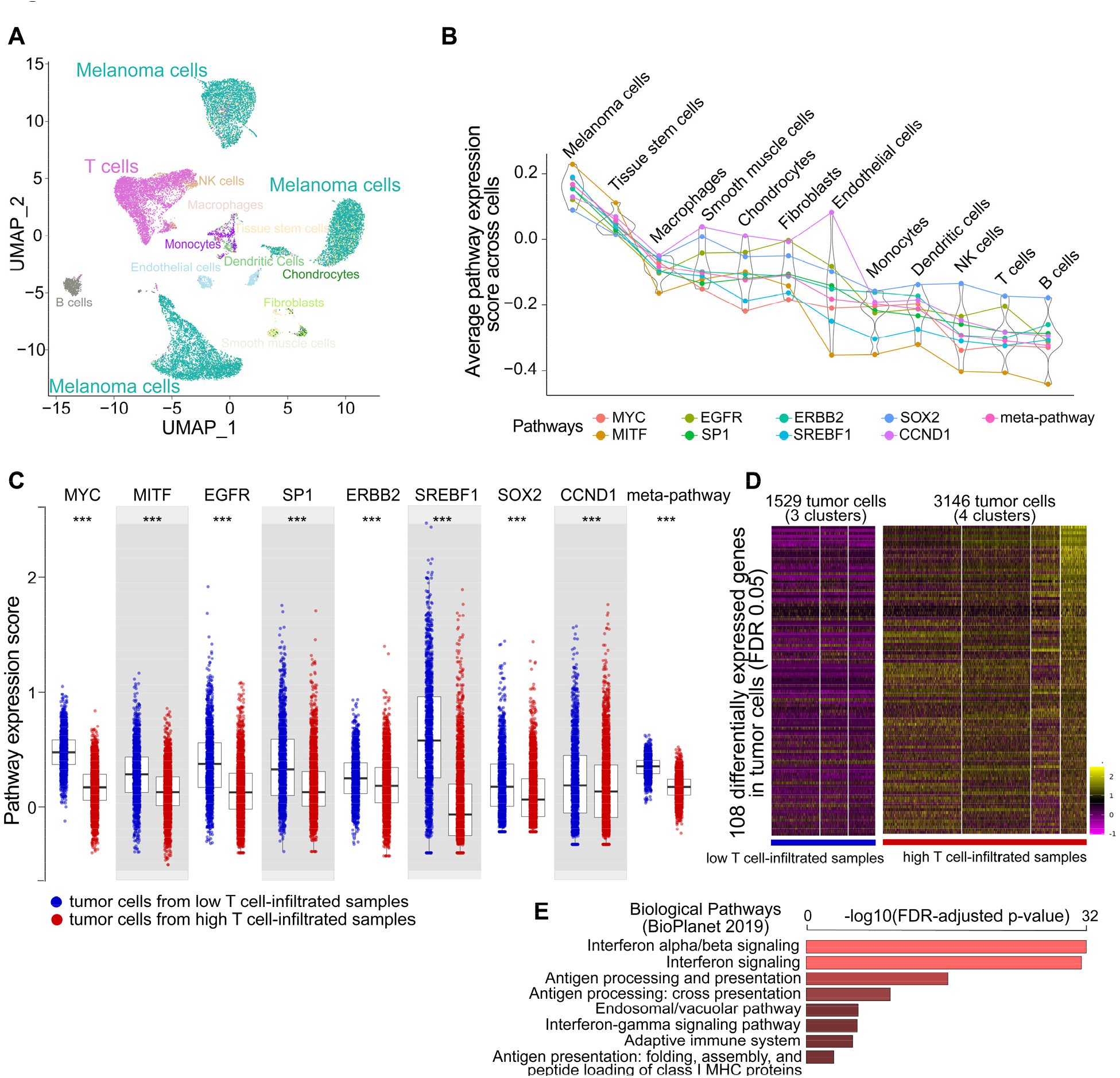
Single cell analysis of immunosuppressive oncogenic pathways in acral melanoma. Data from *Li et al*. are shown after harmonization. (**A**) Cell populations visualized by UMAP. 25523 cells from eight samples are shown (10X Genomics 5’). One sample was excluded from analysis due to different 10X chemistries. Color denotes cell populations annotated by SingleR followed by manual curation. (**B**) Distribution of pathway scores in the main cell populations. (left to right) cell populations were sorted by median pathway scores higher to lower. Eight out of 11 pathways are shown consisting of five to 22 unique downstream target molecules per pathway (MYC, MITF, EGFR, SP1, ERBB2, SERBP1, SOX2, CCND1). Meta-pathway represents a conglomerate expression score of downstream target molecules from all pathways combined. (**C**) Comparison of tumor cell-expressing pathway scores between low and high T cell-infiltrated tumors. Each data point represents one malignant cell. *n*=1529 and 3146 malignant cells shown from low and high T cell-infiltrated tumors, respectively. (**D**) Differentially expressed genes (DEGs) in tumor cells comparing low vs high T cell-infiltrated samples. 108 genes at FDR-adjusted *P*<0.05 shown. Cells are on the column, genes on the row. (**E**) Biological pathway enrichment (BioPlanet2019) on the tumor cell DEGs shown in **D**. P-values were computed by two-sided Two-Sample *t*-test in **C**, two-sided Wilcoxon test in **D**, hypergeometric test in **E**. Denotation: * P <0.05; ** P <0.01; *** P <0.001 after FDR adjustment for multiple comparisons. NK = neutral killer cells.

### Common pathways are upregulated in ICI-resistant tumor cohorts

Expression of the 11 oncogenic pathways associated with a non-T cell-inflamed phenotype was increased within the AM tumors of UPMC patients who had no clinical benefit to ICI (**Fig. 4A**). Given the paucity of harmonized expression and ICI-response data in AM, we analyzed these pathways in two larger CM ICI datasets (**Table S2**). Ten of 11 pathways within Riaz *et al*. (MYC, HGF, MITF, VEGF, EGFR, SP1, ERBB2, TFEB, SOX2, and CCND1; **Fig. 4B upper panel**), and eight of 11 pathways within Liu *et al*. (MYC, HGF, VEGF, EGFR, SP1, ERBB2, SREBF1, SOX2; **Fig. 4B bottom panel**) were found to be upregulated in baseline tumors from patients with who did not have clinical benefit to ICI (FDR-adjusted *P*<0.10). For further validation, we compared the expression of these common pathways with both ICI response and the T cell-inflamed gene signature across three additional CM datasets, one triple-negative breast cancer (TNBC) cohort, one head and neck squamous cell carcinoma (HNSCC) cohort, two non-small cell lung cancer (NSCLC) datasets, one renal cell carcinoma (RCC) cohort, and one urothelial carcinoma cohort, where tissue transcriptomics and ICI clinical data were publicly available (**Table S2**). As depicted in **Fig. 4C**, these pathways are consistently upregulated in baseline tumors from patients who did not have clinical benefit to ICI and inversely correlated with the T cell-inflamed gene signature across several cancers.

**Figure 4.**
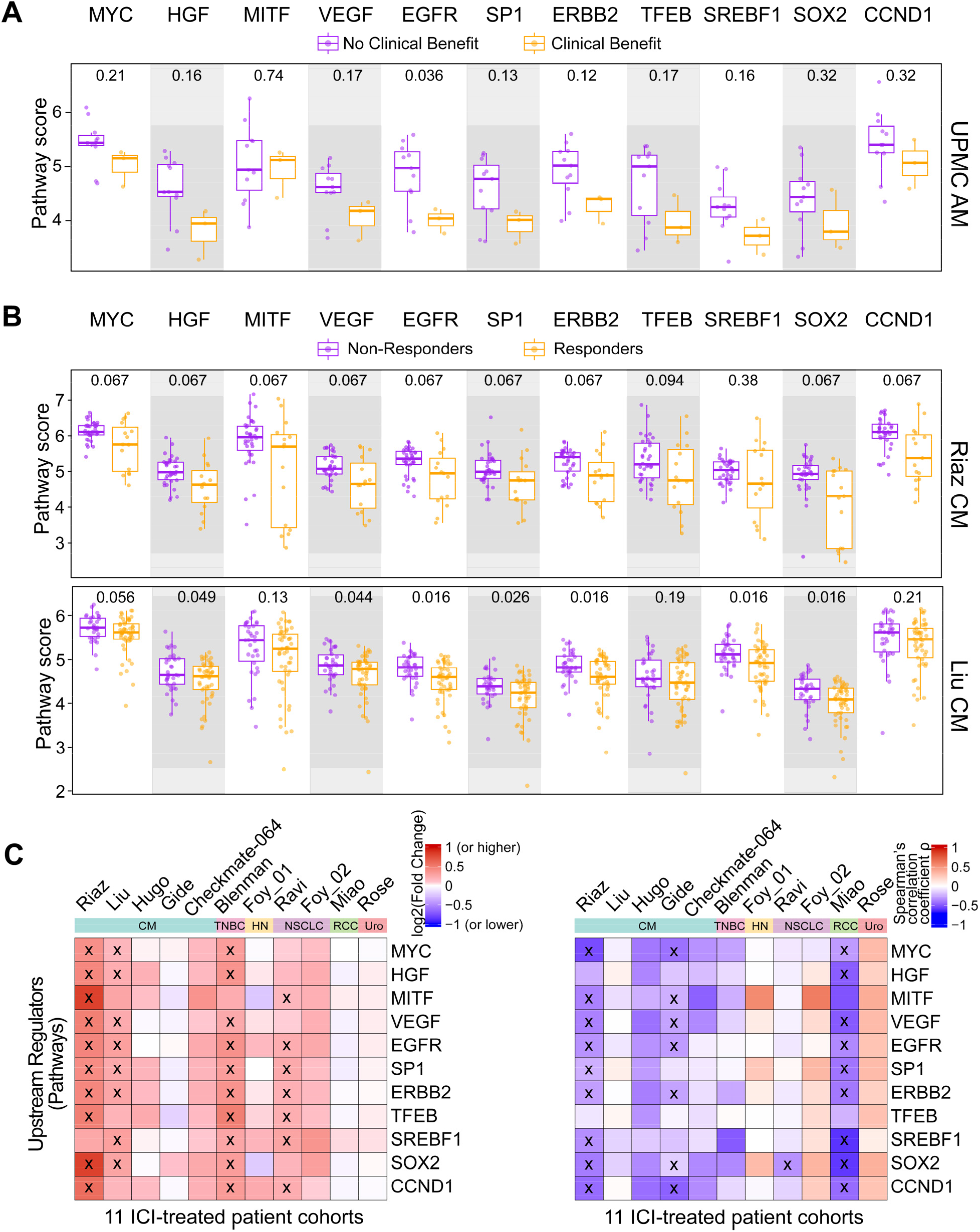
Validation of immunosuppressive oncogenic pathways in ICI-treated cohorts. Baseline (pre-ICI treatment) tumors were used in all analyses. (**A**) Comparison of 11 pathways in acral melanoma (AM) samples from patients who showed clinical benefit (CB) or not (NCB) to ICI. *n*=14 shown, UPMC AM cohort. Patients in NW or MIA AM cohort did not receive ICI, hence not shown. (**B**) Comparison of 11 pathways in cutaneous melanoma (CM) samples from patients who received ICI (Riaz *et al*., and Liu *et al*.). (**C**) Pan-cancer analysis of pathways across ICI-treated patient cohorts. (left panel) The fold change of pathway scores in non-T cell-inflamed relative to T cell-inflamed tumors in each cohort; A label “x” was added to the grids when significant upregulation (log_2_(Fold Change) >0) in NCB vs CB or NR vs R was detected at FDR-adjusted *P*<0.10. (right panel) The correlation between pathway scores and T cell-inflamed gene expression; asterisks were added when significant inverse correlation (Spearman’s ρ <0) was detected at FDR-adjusted *P*<0.10. Eleven ICI datasets with gene expression data and de-identified clinical data publicly available were included in analysis, consisting of five cutaneous melanoma studies (Riaz *et al*., Liu *et al*., Hugo *et al*., Gide *et al*., and Checkmate-064 from Campbell et al.), one triple negative breast cancer study (TNBC; Blenman *et al*.), one head and neck squamous carcinoma study (HN; Foy *et al*.), two non-small cell lung cancer studies (NSCLC; Ravi *et al*., Foy *et al*.), one renal cell carcinoma study (RCC; Miao), and one urothelial carcinoma study (Rose *et al*.). P-values were computed by two-sided Two-Sample *t*-test in **A, B**, and **C upper panel**, Spearman’s correlation in **C bottom panel**.

## Discussion

Tumor-intrinsic mechanisms of immune exclusion have been proposed to be drivers of the non-T cell-inflamed phenotype, leading to resistance to ICI. Whereas Wnt/β-catenin signaling has been identified in CM to drive a lack of CD8^+^ T cell infiltration across human solid tumors, no such analysis has been performed in AM where outcomes to ICI are inferior. By contrasting the non-T cell-inflamed versus T cell-inflamed TME across three independent AM cohorts, we identified 11 pathways as significantly associated with immune exclusion (MYC, HGF, MITF, VEGF, EGFR, SP1, ERBB2, TFEB, SREBF1, SOX2, and CCND1). Analysis of single-cell AM cohorts suggests that these immunosuppressive pathways are primarily expressed from the tumor cells and not other cells of the TME. Moreover, the shared downstream target molecules across these pathways, fatty acid synthase and CXCL8 (*alias* IL-8), may represent high priority targets for therapeutic intervention in non-T cell-inflamed AM.

In considering the 11 immune exclusion pathways identified, all 11 have been previously described to drive innate immune dysfunction or limit CD8^+^ T cell cytotoxicity. Noting that MYC and VEGF are well established mediators of immunosuppression,^13^ knockdown of MITF has been shown to upregulate type I IFN signaling in melanocyte stem cells,^14^ HGF is associated with M2 macrophage polarization, SP1 promotes Treg activity via DcR3, and TFEB has been shown to regulate tumor-associated macrophages in breast cancer.^15,16^ Additionally, SOX2 desensitizes melanoma cells to CD8+ cytotoxicity, CCND1 promotes immunosuppressive features through epithelial-mesenchymal transition (EMT), and SREBF1-mediated fatty acid production is associated with M2 polarization and ICI resistance.^17–19^ Though ERBB1/EGFR-mutant NSCLC is a known predictor of ICI resistance, the role of ERBB2/HER2 in modulating the immune milieu of the TME has been rarely studied outside the context of HER2-directed therapies. Prior studies have, however, associated HER2-mediated fatty acid production and pro-tumor cytokine secretion (e.g. CXCL8) with tumor progression and immune dysfunction in breast cancer models.^20,21^

While functional validation of specific pathways is a future priority, we observe that expression of each pathway was associated with suppression of interferon stimulated genes. A deficit of antigen presentation leading to resistance to ICI would be broadly consistent with what has been described for tumor-intrinsic mechanisms of immune exclusion in other contexts.^7^ For example, STK11 mutation strongly predicts lack of response to ICI in lung cancer where epigenetic silencing of STING is observed.^22^ Similarly, activation of MYC is associated with immunosuppressive tumor hypoxia as well as lack of response to ICI.^11^

Both fatty acid synthase and CXCL8 (*alias* IL-8) were found to be downstream target genes across the 11 pathways in our study. Notably, inhibitors of both the fatty acid synthase and IL-8 axes have entered clinical trials in various solid tumors in combination with ICI. Lipid and cholesterol metabolism have been linked to systemic immunosuppression and reduced rates of response to ICI. Specifically, FASN^high^ ovarian cancer cells have been shown to inhibit tumor-infiltrating dendritic cells via lipid accumulation, and FASN expression is associated with poor ICI efficacy and survival in bladder cancer.^23,24^

Elevated levels of circulating IL-8 have also been associated with poor outcomes to ICI in refractory advanced solid tumors.^25^ The immune inhibitory functions of IL-8 drive several mechanisms including but not limited to neutrophil extracellular traps and myeloid-derived suppressor cell accumulation, EMT, angiogenesis, and proliferation of cancer stem cells, among others.^26^ Breast cancer cells co-expressing HER2 and HER3 have been found to upregulate IL8 expression with resultant tumor invasion, findings reversed with anti-IL8 antibodies.^27^ Another study involving breast cancer mammospheres showed that increasing levels of IL8 exposure were associated with both EGFR and HER2 signaling pathways and tumor growth; inhibiting CXCR1/2 (IL8 cognate receptor) reversed this activity.^28^ An anti-IL-8 antibody (BMS-986253) in combination with ICI has delivered responses in clinical trials of patients with immunotherapy refractory melanoma, including patients with AM.^29^ Our group has further identified IL-8 as a mediator of resistance to stereotactic body radiotherapy plus immunotherapy combinations. Leveraging these observations in identifying IL-8 as a mediator of resistance, we have launched a phase I/II study of SBRT with nivolumab and BMS-986253 (NCT04572451) which will investigate the safety and efficacy of this combination to overcome the non-T cell-inflamed TME in melanoma.

In conclusion, we have identified an AM specific group of tumor-intrinsic pathways tied to immune exclusion and ICI resistance with validation across several tumor cohorts. Given the lack of effective treatment options for patients with AM, we propose further investigation into targeting these pathways to improve immunotherapy outcomes in AM.

## Declarations

### Ethics approval and consent to participate

The study protocol was approved by The University of Pittsburgh institutional review board (IRB)-approved protocol (Protocol No. 20090109). Participants gave informed consent to participate in the study before taking part. All samples have written-informed patient consent.

### Consent for publication

All authors consent.

### Availability of data and materials

The gene expression data will be deposited to NCBI GEO repository. De-identified clinical data is provided in supplementary tables. Public datasets have been referenced as appropriate (**Supplementary Methods**, section “**Study Cohorts and Datasets”**, and **Table S2**). Other data will be provided upon request from the corresponding authors.

### Competing interests

RB declares PCT/US15/612657 (Cancer Immunotherapy), PCT/US18/36052 (Microbiome Biomarkers for Anti-PD-1/PD-L1 Responsiveness: Diagnostic, Prognostic and Therapeutic Uses Thereof), PCT/US63/055227 (Methods and Compositions for Treating Autoimmune and Allergic Disorders); JJL declares DSMB: Abbvie, Immutep; Scientific Advisory Board: (no stock) 7 Hills, Fstar, Inzen, RefleXion, Xilio (stock) Actym, Alphamab Oncology, Arch Oncology, Kanaph, Mavu, Onc.AI, Pyxis, Tempest; Consultancy with compensation: Abbvie, Alnylam, Avillion, Bayer, Bristol-Myers Squibb, Checkmate, Codiak, Crown, Day One, Eisai, EMD Serono, Flame, Genentech, Gilead, HotSpot, Kadmon, KSQ, Janssen, Ikena, Immunocore, Incyte, Macrogenics, Merck, Mersana, Nektar, Novartis, Pfizer, Regeneron, Ribon, Rubius, Silicon, Synlogic, Synthekine, TRex, Werewolf, Xencor; Research Support: (all to institution for clinical trials unless noted) AbbVie, Agios (IIT), Astellas, Astrazeneca, Bristol-Myers Squibb (IIT & industry), Corvus, Day One, EMD Serono, Fstar, Genmab, Ikena, Immatics, Incyte, Kadmon, KAHR, Macrogenics, Merck, Moderna, Nektar, Next Cure, Numab, Pfizer (IIT & industry) Replimmune, Rubius, Scholar Rock, Synlogic, Takeda, Trishula, Tizona, Xencor; Patents: (both provisional) Serial #15/612,657 (Cancer Immunotherapy), PCT/US18/36052 (Microbiome Biomarkers for Anti-PD-1/PD-L1 Responsiveness: Diagnostic, Prognostic and Therapeutic Uses Thereof). P.C.L. declares equity interest in Amgen. D.D. declares grants/research support (NIH/NCI and Checkmate Pharmaceuticals) and consulting (Checkmate Pharmaceuticals) during the conduct of the study. D.D. also reports grants/research support (Arcus, CellSight Technologies, Immunocore, Merck Sharp & Dohme, Tesaro/GSK), consulting [Clinical Care Options (CCO), Finch Therapeutics, Gerson Lehrman Group (GLG), Medical Learning Group (MLG), Xilio Therapeutics], speakers’ bureau (Castle Biosciences) and pending provisional patents related to gut microbial signatures of response and toxicity to immune checkpoint blockade (US Patent 63/124,231 and US Patent 63/208,719) outside the submitted work. J.M.K. declares grants/research support (Bristol-Myers Squibb, Amgen Inc.) and consulting (Bristol-Myers Squibb, Checkmate Pharmaceuticals, Novartis, Amgen Inc., Checkmate, Castle Biosciences, Inc., Immunocore LLC, Iovance, Novartis.) outside the submitted work. H.M.Z. declares grants/research support (NIH/NCI and Checkmate Pharmaceuticals) and consulting (Checkmate Pharmaceuticals) during the conduct of the study, grants/research support (NIH/NCI, Bristol-Myers Squibb and GlaxoSmithKline), personal fees (GlaxoSmithKline and Vedanta) and pending provisional patents related to gut microbial signatures of response and toxicity to immune checkpoint blockade (US Patent 63/124,231 and US Patent 63/208,719) outside the submitted work. Correspondence and requests for materials should be addressed to J.J.L. and R.B. The remaining authors declare no competing interests.

### Funding

This work was supported by National Institutes of Health (NIH) Grant R01DE031729 (R.B., J.J.L.), P50CA097190 (R.B.), UM1CA186690 (J.J.L.), P50CA254865 (R.B., J.J.L., D.D., J.M.K, H.M.Z.), in part by National Cancer Institute through the UPMC Hillman Cancer Center CCSG award (P30CA047904), and in part by The University of Pittsburgh Center for Research Computing through the resources provided, specifically the HTC high-performance computing cluster supported by NIH award number S10OD028483.

### Authors’ contributions

R.B. conceived and designed the overall study and oversees computational data analysis and experimental design. J.J.L. designed study and oversees clinical review. R.B. and J.J.L., editorial oversight. R.C.A. wrote IRB, designed methods, and performed clinical and computational analyses. S.N. coordinated tumor sample collection and processing, A.L. performed pathology annotation, M.J. optimized the mIFpanels with PhenoImager^TM^ HT, performed staining, and spectrally unmixed the imaging data; M.L. performed tumor microdissection and sequencing library preparation; K.S. performed immunohistochemistry staining; C.S. performed tumor sample collection and de-identified clinical data acquisition; B.I. performed imaging analysis; D.D., Y.G.N., H.M.Z., J.M.K. contributed tumor samples and patient data.R.C.A., J.J.L., and R.B. wrote the manuscript. All authors contributed to the final manuscript.

## Acknowledgments

We are grateful to Shantel Olivares and Pedram Gerami from Northwestern University along with Felicity Newel and Nicholas Hayward from the Melanoma Institute of Australia for providing access to gene expression data from their published studies. The whole transcriptome RNAseq and whole-exome sequencing were performed at UPMC Hillman Cancer Center Cancer Genomics Facility (CGF) and UPMC Genome Center (UGC), respectively. The authors thank F. Mu (U. Pittsburgh) for their technical assistance in software installation and job execution on the HPCs.

## List of abbreviations

TCGA: The Cancer Genome Atlas
PE: paired-end
IgG: immunoglobulin G
IHC: Immunohistochemistry
CPM: counts per million of mapped reads
DEG: Differentially expressed genes
IPA: Ingenuity Pathway Analysis
TMB: tumor mutational burden
OS: overall survival
FDR: false-discovery rate
mIF: multispectral immunofluorescence
WES: whole exome sequencing
RNAseq: RNA sequencing
TIL: tumor infiltrating lymphocyte
AM: acral melanoma
CM: cutaneous melanoma
NW: Northwestern University
MIA: Melanoma Institute of Australia
UPMC: Hillman Cancer Center
NSCLC: non-small cell lung cancer
TNBC: triple-negative breast cancer
HNSCC: head and neck squamous cell carcinoma
RCC: renal cell carcinoma
TME: tumor microenvironment
ICI: immune checkpoint inhibitor
CB: clinical benefit
FFPE: formalin-fixed paraffin embedded
IFN: interferon
SBRT: stereotactic body radiotherapy
HR: hazard ratio
EMT: epithelial-mesenchymal transition.

