## Supplemental Materials for "Identification of tumor-intrinsic drivers of immune exclusion in acral melanoma"

### For

**Running Title:** acral melanoma immune exclusion

**Feature**: Immunotherapy Biomarkers

**Authors**: Ryan C. Augustin,[1,2] Sarah Newman,[1] Aofei Li,[3] Marion Joy,[1,2] Maureen Lyons,[1] Mary Pham,[1] Peter C. Lucas,[1,3] Kate Smith,[1] Cindy Sander,[1] Brian Isett, [1] Diwakar Davar,[1,2] Yana G. Najjar,[1,2] Hassane M. Zarour,[1,4] John M. Kirkwood,[1,2] Jason J. Luke,[1,2*] Riyue Bao[1,2*]

[1]UPMC Hillman Cancer Center, Pittsburgh, PA
[2]University of Pittsburgh, Department of Medicine, Pittsburgh, PA
[3]University of Pittsburgh, Department of Pathology, Pittsburgh, PA
[4]University of Pittsburgh, Department of Immunology, Pittsburgh, PA
* Co-senior and corresponding authors

[*]**Corresponding Authors:**
Jason J. Luke, MD, FACP
Associate Professor of Medicine
UPMC Hillman Cancer Center
5150 Centre Ave. Room 1.27C
Pittsburgh, PA 15232


Riyue Bao, PhD
Associate Professor of Medicine
UPMC Hillman Cancer Center
5150 Centre Ave, Suite 1A, Room 105
Pittsburgh, PA 15232


**Keywords**: acral melanoma, immunotherapy, checkpoint inhibitors, tumor microenvironment, immune exclusion, therapeutic targets, immunogenomics, tumor-intrinsic pathways, T cell-inflamed, non-T cell-inflamed

This document contains the full description of **Methods** for the manuscript, **Supplementary Figures 1 to 3,** and **Title of Supplementary Tables**. The **Supplementary Tables 1 to 7** were supplied as separate spreadsheets**.**

#### Supplementary Methods

##### Human melanoma specimens

Formalin-fixed, paraffin-embedded (FFPE) tumor tissue samples were obtained from the melanoma biospecimen bank of UPMC Hillman Cancer Center (UPMC) (*n*=20; 14 patients were treated with anti-PD1). The study protocol was approved by The University of Pittsburgh institutional review board (IRB)-approved protocol (Protocol No. 20090109). All samples have written-informed patient consent.

##### Clinical data annotation and ICI response

Demographic, histopathologic, treatment, and response variables were collected for each UPMC sample using de-identified records as per our IRB-approved protocol (**Table S1**). For consistency, out of 20 patients in total, 14 patients treated with anti-PD1 checkpoint blockade were included in the response comparisons. The remaining five patients were not included in the response analysis due to treatment with anti-CTLA4 alone, insufficient time for response, or because of no reported recurrence with adjuvant ICI treatment only. Of the 15 patients treated with anti-PD1 therapy, four were treated with anti-PD1 in the first-line metastatic setting following adjuvant therapy, two were treated with adjuvant anti-PD1 alone (with disease progression), one was treated with first-line dual CTLA4/PD1 blockade, two were treated with first-line anti-CTLA4 with subsequent anti-PD1, and six were treated with first-line anti-PD1 alone. Clinical benefit to PD1 inhibition was defined as greater than six months of progression-free survival (PFS).[1]

**Study Cohorts and Datasets**

We collected 16 cohorts of multi-omics data from 892 clinically annotated human specimens for the integrative analysis in this study (**Table S2**). We used 109 acral melanoma samples for discovery and validation, and additionally utilized 247 CM samples for comparative analysis of melanoma TME between the two subtypes. All data were from baseline tumors unless otherwise noted.

Datasets include: (1) acral melanoma specimens in the UPMC melanoma biobank (*n*=20; referred to as, UPMC AM cohort). RNAseq, whole-exome sequencing (WES), IHC, and multispectral immunofluorescence data were generated from the 20 AM tumors in this study; (2) acral melanoma RNAseq and clinical data from Northwestern University (*n*=22; NW AM cohort[2]), (3) acral melanoma RNAseq, clinical data, and whole-genome sequencing (WGS)-detected somatic mutations from Melanoma Institute of Australia (*n*=67; MIA AM cohort[3]), (4) AM single-cell(sc) RNAseq dataset (n=8; AM scRNAseq cohort[4]), (5) second AM single-cell(sc) RNAseq dataset (n=6),[5] (6) CM RNAseq gene expression and ICI response data (Riaz, *n*=30),[6] (7) second CM RNAseq set with expression and ICI response data (Liu, n=84),[7] (8) third CM RNAseq set with expression and ICI response data (Hugo, n=18),[8] (9) fourth CM RNAseq set with expression and ICI response data (Gide, n=73),[9] (10) fifth CM RNAseq set with expression and ICI response data (CheckMate-064, n=42),[10] (11) triple-negative breast cancer (TNBC) with RNAseq and neoadjuvant ICI response data (Blenman, n=50),[11] (12) head and neck squamous cell carcinoma (HNSCC) with targeted RNAseq and ICI response data (Foy HNSCC, n=102),[12] (13) non-small cell lung (NSCLC) cohort with RNAseq and ICI response data (Ravi, n=152),[13] (14) second NSCLC cohort with targeted RNAseq and ICI response data (Foy NSCLC, n=82),[12] (15) renal cell carcinoma (RCC) cohort with RNAseq and ICI data (Miao, n=33),[14] and (16) urothelial carcinoma cohort with RNAseq and ICI data (Rose, n=103).[15]

Clinical variables were used as appropriate to filter the publicly available, non-acral, ICI-treated tumors for response comparison and T cell-inflamed gene expression correlation among the 11 common pathways. In general, all datasets were filtered for pre-treatment samples, CM datasets were filtered for cutaneous/skin samples only, stable disease was excluded from the response variable in the Foy HNSCC and NSCLC datasets, stage I patients were excluded from Foy NSCLC, and the TNBC cohort was the only dataset that contained an IHC $PDL1^{+}$ variable for response evaluation in the perioperative setting.

**Whole transcriptome RNAseq library preparation and sequencing**

The nuclear extraction, macrodissection, library preparation, and sequencing were performed at UPMC Hillman Cancer Center Cancer Genomics Facility (GCF). One to six, 10-micron slides were deparaffinized and the area of interest was macrodissected prior to RNA extraction. Slides were washed 3 times in 100% xylene for 5 minutes each followed by air drying prior to dissection. For each sample, the FFPE blank slides were aligned with the stained H&E with the area of interest outlined. The area of interest was scraped using a sterile, #10 scalpel and nuclease-free water. Tissue was then placed into nuclease-free, low retention, 2 ml Eppendorf tube. Samples were additionally washed 2 times with 500 µL of Xylene followed by 500 µL of 100% ethanol and 500 µL of 75% ethanol. The dissected, deparaffinized tissue was air dried prior to adding 240 µL of Qiagen Buffer PKD (Qiagen, Germantown, MD, catalog #: 1034963), and 10 µL of QIAGEN Proteinase K (catalog #19131). RNA was extracted using the RNeasy FFPE kit (Qiagen, Germantown, MD, catalog #:73504) according to the manufacturer's instructions. RNA concentration was determined using the High Sensitivity RNA Qubit kit (ThermoFisher Scientific, Pittsburgh, PA, catalog#: Q32855) and the RNA quality was determined using the NanoDrop and the Agilent Pico RNA Bioanalyzer kit (Agilent, Santa Clara, CA, catalog#:5067-1513). 100 ng of total RNA underwent library prep using the TruSeq® RNA Library Prep for Enrichment (Illumina, San Diego, CA, catalog#:20020189) assay for library construction according to the manufacturer's instructions (Illumina TruSeq RNA Exome reference guide: document # 1000000039582 v01, March 2018). As per the protocol recommendations for FFPE RNA, the samples were not fragmented prior to cDNA synthesis. Samples underwent 15 cycles of pre-hybridization PCR using the following protocol: 98°C/30sec, then 15 cycles of: 98°C/10sec, 60°C/30sec, 72°C/30sec, then 72°C/5min, 4°C Hold. The concentration and size distribution of the pre-hybridization PCR product was determined using the double stranded High Sensitivity DNA Qubit kit (ThermoFisher Scientific, Pittsburgh, PA, catalog#: 32854) and the Agilent High sensitivity DNA Bioanalyzer kit (Agilent, Santa Clara, CA, catalog#: 5067-4626). Samples had an average distribution of approximately 260 base pairs. 200 ng of cDNA per sample was used for individual hybridizations (Illumina TruSeq® RNA Enrichment kit, catalog #: 20020490) and the Illumina exome panel (Illumina catalog #: 20020183). Final library amplification of 10 cycles using the following protocol: 98°C-30sec, then 10 cycles of 98°C-10sec, 60°C-30sec, 72°C-30sec, 72°C-5min, 10°C Hold. Upon the final library PCR clean-up, each library was individually interrogated using the double stranded High Sensitivity DNA Qubit kit (ThermoFisher Scientific, Pittsburgh, PA, catalog#: 32854) and the Agilent High sensitivity DNA Bioanalyzer kit (Agilent, Santa Clara, CA, catalog#: 5067-4626). Prior to sequencing, the concentration of each sample was confirmed using the double stranded high sensitivity DNA Qubit kit (ThermoFisher Scientific, Pittsburgh, PA, catalog#: 32854). Individual samples were then diluted down to 4 nM, following the Illumina NextSeq Denature and dilution guide (Illumina Document # 15048776 v18). The pooled library was denatured and diluted to 1.5 pM final library concentration. Sequencing was in batches of sixteen on the Illumina NextSeq 500/550 using a 150-cycle high output kit. The sequencing run was set on the NextSeq using the Illumina NextSeq Local Run Manager with the following parameters: paired end, R1 76 base pair X R2 76 base pair, with single 8 base pair index.

**Whole exome library preparation and sequencing**

The nuclear extraction, macrodissection, and library preparation were performed at

UPMC Hillman Cancer Center Cancer Genomics Facility (GCF); and sequencing was performed at UPMC Genome Center (UGC). One to seven, 10-micron slides were deparaffinized and the area of interest was macrodissected prior to DNA extraction. Slides were washed 3 times in 100% xylene for 5 minutes each followed by air drying prior to dissection. For each sample, the FFPE blank slides were aligned with the stained H&E with the area of interest outlined. The area of interest was scraped using a sterile, #10 scalpel and nuclease-free water. Tissue was then placed into nuclease-free, low retention, 2 ml Eppendorf tube. Samples were additionally washed 2 times with 500 µL of Xylene followed by 500 µL of 100 % ethanol and 500 µL of 75% ethanol. The dissected, deparaffinized tissue was air dried prior to adding 300 µL of Qiagen Buffer ATL (Qiagen, Germantown, MD, catalog #: 939011), and 100 µL of QIAGEN Proteinase K (catalog #19131). DNA was extracted using the QIAamp DNA FFPE Tissue Kit (Qiagen, Germantown, MD, catalog #: 56404). Samples were incubated in a shaking heat block at 56 °C for >24 hours depending on the size of the area excised (Lysis time varied from 1 to 5 days). DNA extraction was performed according to the manufacturer’s instructions with the exception of using an excess volume of Buffer ATL and Proteinase K. An excess of both was used as the lysis time was > 24 hours. gDNA concentration was determined using the double stranded DNA High Sensitivity Qubit kit (ThermoFisher Scientific, Pittsburgh, PA, catalog#: 32854) and the gDNA quality was determined using the NanoDrop and the Agilent High sensitivity DNA Bioanalyzer kit (Agilent, Santa Clara, CA, catalog#: 5067-4626). 200 ng of gDNA underwent exome library prep using the Agilent Sure Select XT library kit (Agilent, Santa Clara, CA, catalog#: G9611B) assay for library construction according to the manufacturer’s instructions (SureSelectXT Target Enrichment System for Illumina Paired-End Multiplexed Sequencing Library protocol (Agilent, Santa Clara, CA, Version C3, September 2019). Specifically, 0.25ng/ul (50 ul low TE) ds gDNA was pipetted into Covaris microtube AFA Fiber pre-slit snap cap (6x16) tubes (Covaris, Woburn, MA, catalog#: 520045). Samples were sheared using the Covaris S-1 (Covaris, Woburn, MA) with the following settings (Duty Cycle:

10%, Intensity: 5, Cycles per burst: 200, Time: 8 cycles of 40 seconds for a total of 320 seconds, Set Mode: Frequency Sweeping, Temperature: 4°C to 7°C). Achieving an average fragment size of approximately 150 base pair. The Agilent High sensitivity DNA Bioanalyzer kit (Agilent, Santa Clara, CA, catalog#: 5067-4626) was used to confirm the fragment size of each sample. Once confirmed, DNA was end repaired, A's added, ligated, and underwent pre-hybridization PCR (98°C 2 minutes, 12 cycles of: 98°C 30 seconds, 65°C 30 second, 72°C 1 minute, 72°C 1 minute). The concentration and quality of the pre-hybridization PCR product was determined using the double stranded High Sensitivity Qubit kit (ThermoFisher Scientific, Pittsburgh, PA, catalog#: 32854) and the Agilent High sensitivity DNA Bioanalyzer kit (Agilent, Santa Clara, CA, catalog#: 5067-4626). Samples had an average distribution between 225 to 275 base pairs. Exome hybridization was performed using the Agilent Human all exon V6 baits (Agilent, Santa Clara, CA, catalog#: 5190-8863) using 750 ng of each sample. All samples underwent individual, 24-hour hybridization to the exome baits. Exome capture and post hybridization library prep was performed using Agilent's SureSelect XT hybridization and blocking reagents (Agilent, Santa Clara, CA, catalog#: 930672) and 50 ul of MyOne Streptavidin T1 beads (ThermoFisher Scientific, Pittsburgh, PA, catalog #:65602). Final library amplification of 11 cycles added the individual index adaptors to the captured libraries. PCR conditions of (98°C 2 min; 98°C 30 sec., 57°C 30 sec., 72°C 1 min (repeat for 11 cycles) and 72°C for 1 min). For all clean-up steps, we used AMPure XP beads (Beckman Coulter catalog# A63881, Indianapolis, IN 46268) and 70% ethanol. Beads were incubated for 5 minutes, washed twice with 70% ethanol, and dried on a 37°C heat block for less than 5 minutes prior to elution. Final library concentration and average library size was determined using the double stranded High Sensitivity Qubit kit (ThermoFisher Scientific, Pittsburgh, PA, catalog#: 32854) and the Agilent High sensitivity DNA Bioanalyzer kit (Agilent, Santa Clara, CA, catalog#: 5067-4626). Final libraries were transferred to the UPMC Genome Center for sequencing. Prior to sequencing, the concentration of each sample was confirmed using the double stranded high sensitivity DNA

Qubit kit (ThermoFisher Scientific, Pittsburgh, PA, catalog#: 32854). Individual samples were then diluted down to 4 nM, following the Illumina NextSeq Denature and dilution guide (Illumina Document # 15048776 v18). The pooled library was denatured and diluted to 1.5 pM final library concentration. All WES samples were included in the same run for sequencing an Illumina NovaSeq 6000 instrument using a 200-cycle high output kit to generate paired end, R1 100 base pair X R2 100 base pair, with single 8 base pair index.

**Identification of T cell-inflamed and non-T cell-inflamed tumor groups**

A predefined and validated 160-gene T cell-inflamed gene signature was utilized to categorize tumors into T cell-inflamed, intermediate, or non-T cell-inflamed using consensus clustering methods following previous protocols.[16–18] Briefly, an expression matrix consisting of the 160 genes from the T cell-inflamed signature was subset from the TMM-normalized and $log_2$-transformed RNAseq gene expression quantification matrix, and was used to cluster tumors into three clusters using hierarchical clustering with Euclidean distance and Ward.D2 linkage using the ComplexHeatmap package (v2.10.0). Tumors were then assigned to each of the three immune groups based on high, low, or intermediate expression of the T cell-inflamed signature. This signature has been previously validated, showing strong correlation with established markers of T cell inflammation and other inflammatory gene signatures.[18] The 160-gene T cell-inflamed gene signature was built upon a 13-gene signature that we originally established in metastatic melanoma[19] and further on expanded to be broadly applicable across 31 human solid tumors from The Cancer Genome Atlas (TCGA), as demonstrated in our previous studies.[16–18] To summarize the methodology, for each specific cancer type, the normalized and log2-transformed RNAseq gene expression matrix underwent unsupervised hierarchical clustering using K equal to 12 and Euclidean distance. The resulting clusters that encompassed the original 13 T-cell gene signature were identified. Subsequently, a set of 160 genes consistently co-clustered with these 13 genes was extracted, hereafter referred to as the T cell-inflamed gene signature.[16–18]

#### RNAseq gene expression quantification

For UPMC AM, NW AM, and TNBC cohorts, we started with the raw FastQ files (76 bp PE reads for both cohorts) and processed all data to generate count matrices in house using kallisto (v0.48.0),[20] same as our previous work.[18] In brief, after quality control, reads were pseudoaligned via kallisto (v0.48.0) with human reference transcriptome (GRCh38) and Gencode annotation (v28), summarized into gene level by tximport (v1.22.0). From here and for other RNAseq datasets, raw count matrices were directly downloaded (sources above), followed by TMM normalization. All expression values were $\log_2$-transformed before statistical analysis.

#### Differential gene expression detection and pathway activation prediction

For each RNAseq dataset, after removing genes with low expression (defined as CPM (counts per million of mapped reads) ≤ 1), differentially expressed genes (DEGs) were calculated by contrasting the non-T cell-inflamed groups against the T cell-inflamed groups using Linear Models for Microarray and RNA-Seq Data (limma) voom algorithm with precision weights (v3.50.3).[21] Significant DEGs were filtered by $P < 0.05$ and fold-change ≥ 1.5 or ≤ − 1.5. Upstream transcriptional regulators and change of direction (activation or inhibition) as a result of target molecules (encoded by DEGs) was predicted using Ingenuity Pathway Analysis (IPA®) (QIAGEN Inc., Germany) causal network analysis with the curated Ingenuity Knowledge Base (accessed 8/2022 – 4/2023). Transcriptional programs activated in non-T cell-inflamed relative to T cell-inflamed tumors were filtered at overlap $P<0.05$ (measuring the enrichment of target molecules in the dataset) and z-score ≥ 1.95 (measuring the predicted activation level of the pathways).

#### T cell-inflamed gene expression and pathway score calculation

For each tumor, a T cell-inflamed score was computed as the mean expression of the 160 genes involved in the signature after scaling and centering across all tumor samples.[16] In addition, for each pathway identified in this study (MYC, HGF, MITF, VEGF, EGFR, EGFR, SP1, ERBB2, TFEB, SREBF1, SOX2, and CCND1), the expression score of a pathway was defined by the mean expression of all target molecules from this pathway. These expression scores were then used to correlate with the T cell-inflamed gene expression across all tumors by Spearman's correlation. Pathway activation scores were calculated for each individual tumor sample, requiring at least 50% of the cancer-specific target molecules to be upregulated (relative to its median expression across all samples from an individual cohort) and compared between the non-T-cell-inflamed tumor group relative to inflamed. For pathways where the total number of target molecules is less than 10, at least five molecules are required to be upregulated in order to conclude this pathway is activated in a tumor.

**Somatic mutation detection and TMB quantification**

Somatic mutations in AM tumors were detected following GATK best practice guidelines. After QC, paired-end reads were aligned to human reference genome GRCh38 using BWA MEM (v0.7.17) with mapping quality ≥30, followed by duplicate removal and base quality score recalibration (BQSR). The refined read alignment was used for detecting putative somatic variants using GATK4-MuTect2 (v4.3.0.0) with tumor-only mode and a panel of normal (PoN) built by 30 WES germline samples sequenced at the same facility. GATK4-MuTect2 employs intrinsic quality and background noise filters to remove false positives. We further removed low-quality somatic variant calls and potential germline leaking by (1) eliminating strand orientation bias, (2) eliminating OxoG artifacts, (3) removing variants present in gnomAD exome or genome database (v2.1.1) at allele frequency ≥ 0.0001, and (4) requiring at least 8 support reads to make a confident call on putative somatic variants. Somatic variants that passed all filters were carried on for annotation by GATK4-Funcotator. Total TMB was defined as the total number of non-synonymous somatic mutations (NSSMs), those that were predicted to alter protein sequence in tumor (insertions/deletions, missense/nonsense/stop-gain mutations, and those that modify splicing sites).

**Single-cell RNAseq analysis**

For the two scRNAseq AM cohorts, data were harmonized in this study using the same bioinformatics pipelines. Raw FastQ Files were downloaded from the public NCBI SRA repository (Li *et al.*, PRJNA784961; Zhang *et al.*, PRJNA862451), and processed by Cell Ranger (v6; 10x Genomics) to generate the single-cell Unique Molecular Identifier (UMI) count matrix with proper denoising and filtering. Cells with <200 or >5000 genes expressed, or >15% mitochondria gene expression were excluded. Data was normalized by scaling expression values to 10,000 counts per cell, followed by log transformation. After selecting the first 10 principal components using Principal Component Analysis (PCA), a neighborhood graph was computed based on the PCA representation of the data and visualized by Uniform Manifold Approximation and Projection (UMAP) dimensionality reduction technique with Leiden clustering. Cell populations were annotated using SingleR (v1.10.0) with reference markers from HumanPrimaryCellAtlasData. Malignant (tumor) cells were identified by CopyKat (v1.1.0) with immune cells and fibroblasts specified as the normal cells. Samples that are treatment naïve, and passed the QC step in CopyKat were used for further statistical comparison of tumor cell-expressing pathway scores in low vs high T cell-infiltrated tumors.

**H&E and CD8 immunohistochemistry staining**

The hematoxylin and eosin (H&E) and CD8 IHC staining on UPMC AM specimens ($n$=17) was performed at UPMC Developmental Laboratory. Slides were cut at 4μm then baked for one hour at 60 degrees Celsius. The slides were cooled to room temperature then deparaffinized and hydrated in diH20. Slides were stained using Hematoxylin 560 MX (3801576,

Eosin Phloxine 515 (3801606), Define MX-aq (3803598) and Blue Buffer 8 (3802918) (all catalog numbers from Leica Biosystems). The predilute CD8 (Leica Biosystems, PA0183) was stained on the Bond III instrument (Leica Biosystems) for 8 min. Prior to antibody placement, the antigen retrieval used was the ER2 (pH 8.9-9.1, Leica Biosystems, AR9640) for 20 minutes at 95 deg. Followed by the Bond Polymer Refine Detection Kit-DAB (Leica Biosystems, DS9800).

**CD8 IHC quantification**

IHC whole slide images (WSI; 40X) were imported into QuPath (v0.4.3) with default settings. Cells were segmented and DAB+ cells were detected, with cell segmentation parameters medianRadiusMircons=1, sigmaMicrons=1.5, Hematoxylin threshold = 0.05; positive cell detection using DAB+ threshold = 0.2 nuclear DAB mean. Tumor regions were annotated on the H&E section of each tumor. By aligning these cell detections to H&E tumor area, IHC DAB+ cell density was measured inside and outside tumor region of registered cores as the percent of DAB+ out of all cells detected for each corresponding IHC WSI.

**Multispectral immunofluorescence Vectra staining and imaging**

Multispectral immunofluorescence staining was performed on 4um tumor FFPE sections: CD3 (cat# 85061S), CD8 (cat# ACI3160A), PD1 (cat# ab137132), SOX10 (cat# ACI3099A), and DAPI. Automated staining of tissues was performed on the Leica Bond RX. For staining, Akoya Bioscience's Opal 6-Plex Manual Detection Kit was used according to the manufacturer's instructions (cat# NEL861001KT). Imaging was performed at 20X on the Vectra Polaris 3.0. InForm® (v2.4.6) and Phenochart™ (v1.0) (Akoya Biosciences, Inc.) analysis software was used for whole slide scanning and regions of interest (ROI) spectral unmixing. Multi-channel composite TIFF files were exported for further data analysis.

**Multispectral immunofluorescence Vectra image analysis**

ROIs from the same WSI were stitched together to constitute the full tumor region at 20X resolution. Cell segmentation was performed on DAPI channel using a Stardist model that we trained using 20K manually annotated cells across three platforms. Marker positive/negative cells were classified using a machine learning approach. Briefly, a small number of positive/negative cells (30–50 per class per ROI) were manually selected as the training set to build a Random Forests model, which was then used to predict all remaining cells. Tumor/peritumoral/stromal compartment annotation was performed using pixel classification by SOX10 intensity signals. Measurement matrices consisting of centroid position (x,y), per-channel intensity, and class label of phenotyped cells were exported and further processed in R (v4.1.2). $CD3^+CD8^+PD1^+$ T cell density per 10000 cells was computed within each compartment.

**Survival analysis**

Kaplan-Meier (KM) estimator with log-rank test (nonparametric) was used to compare the overall survival (OS) trend between patients stratified by the $CD3^+CD8^+PD1^+$ T cell density, or tumor groups (T cell-inflamed, non-T cell-inflamed, and intermediate), in the UPMC AM cohort using R package survminer (v0.4.9). Cox proportional hazard (PH) univariable or multivariable regression models were used to test the association between T cell-inflamed gene expression as a continuous variable and OS (with age, sex, TMB as covariates). Right censoring was used.

**Statistical analysis**

The t-test was used to compare mean expression values between tumor groups (response and T cell-inflamed variables). Differential gene expression analysis between groups was performed using empirical Bayes regression models in limma voom with precision weights. Spearman’s correlation was used for measuring statistical dependence between normalized and

$log_2$-transformed expression level of different genes, and between gene expression of the T cell-inflamed signature and pathways. *P*<0.10 was considered statistically significant. Benjamini-Hochberg (BH) Procedure was used to adjust multiple comparisons when appropriate. All tests are two-sided unless otherwise noted. Statistical analysis was performed using R (v4.1.2) and Bioconductor (release 3.14).

### Supplementary Figures

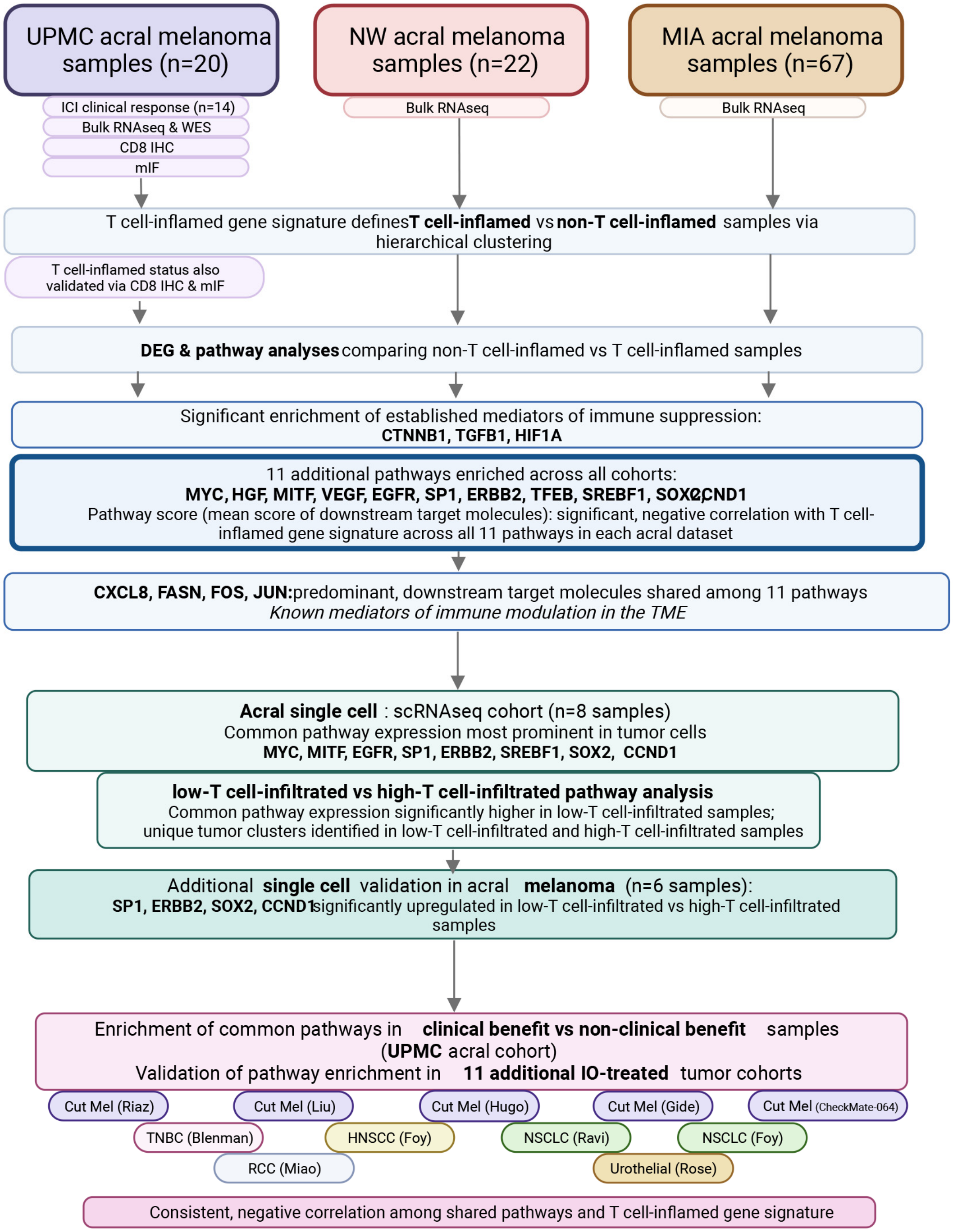


**Supplementary Figure 1. Workflow of cohorts analyzed, experiments performed, and key findings.**

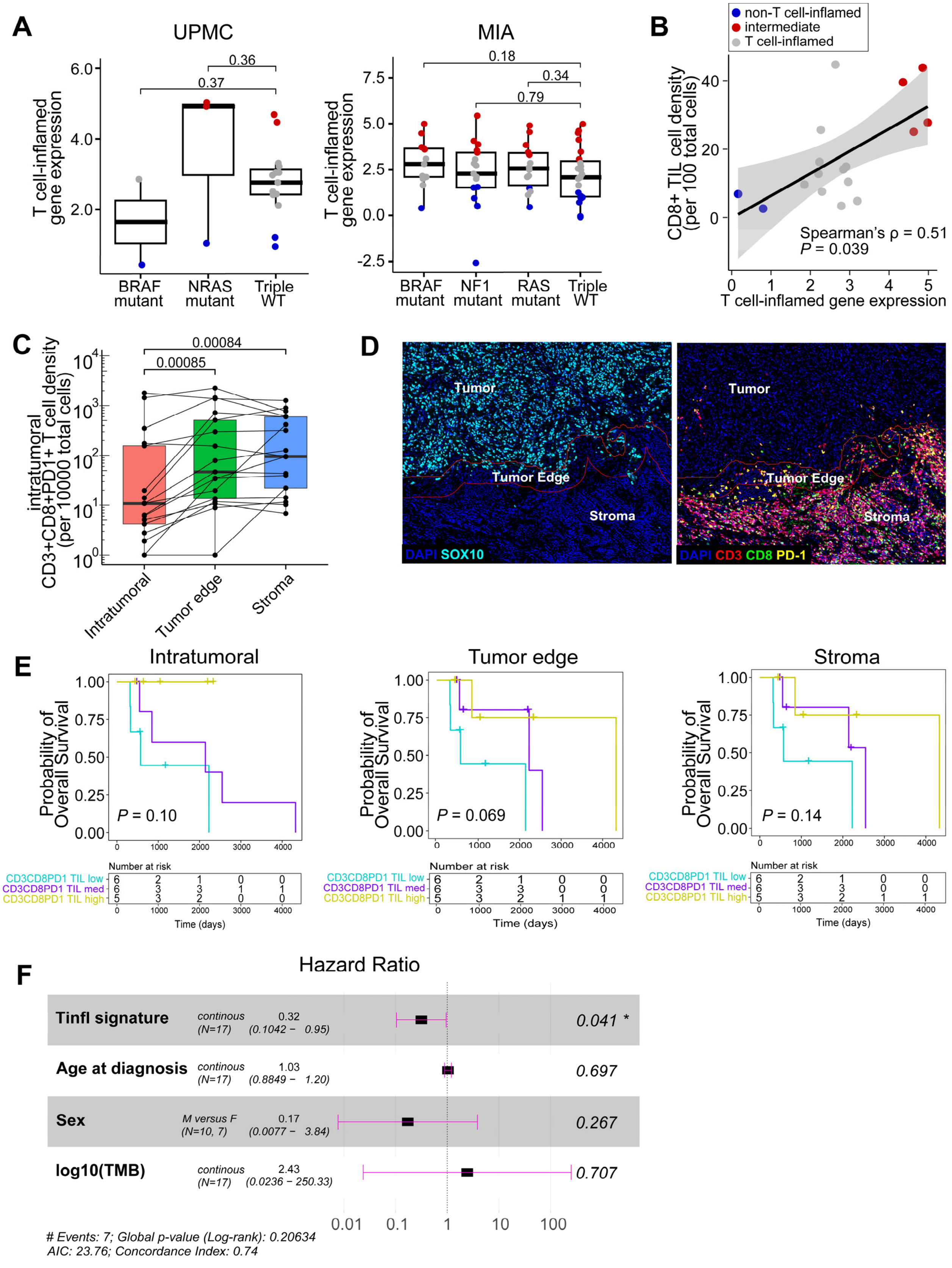


**Supplementary Figure 2. Immune infiltration and other correlatives in the acral melanoma**

**TME.** (**A**) Comparison of T cell-inflamed gene expression signature between typical melanoma driver mutations (UPMC, MIA). NW cohort does not have mutation data available. No tumors carry an NF1 mutation in the UPMC cohort. Each data point represents one tumor sample. (**B-F**) *n*=17 shown with tumor specimens available for imaging experiments from the UPMC cohort. (**B**) Correlation between T cell-inflamed gene expression signature and $CD8^+$ tumor infiltrating lymphocyte (TIL) cell density measured by IHC. (**C**) Distribution of $CD3^+CD8^+PD1^+$ T cell density in intratumoral, tumor edge, or stromal space via multispectral imaging. (**D**) Representative immunofluorescence images showing intratumoral, tumor edge, and stroma compartments. (**E**) Kaplan-Meier survival curves of patients' overall survival (OS) stratified by $CD3^+CD8^+PD1^+$ T cell infiltrate low/med/high split by a third. (left to right) cell density in intratumoral, tumor edge, or stromal space illustrated in **C** and **D.** (**F**) Cox PH multivariable model of T cell-inflamed gene expression signature in association with OS, with age, sex, and TMB as covariates. P-values were computed by two-sided Two-Sample *t*-test in **A**, Spearman's correlation in **B**, two-sided Wilcoxon test in **C**, log-rank test in **E**, and Wald test in **F**.

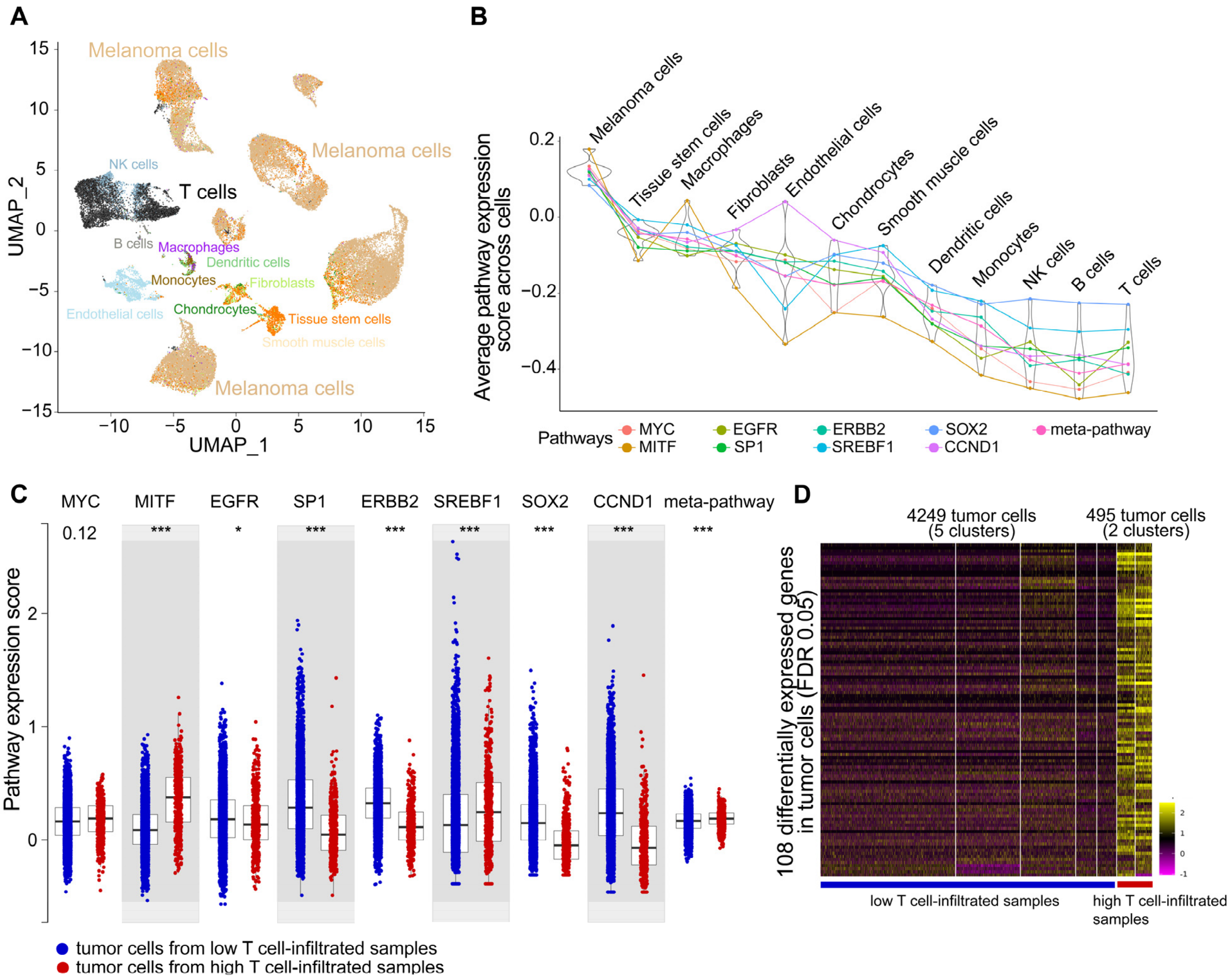


**Supplementary Figure 3**. **Single cell analysis of immunosuppressive oncogenic pathways in acral melanoma.** Data from *Zhang et al.* are shown after harmonization. (**A**) Cell populations visualized by UMAP. 42558 cells from six samples are shown (10X Genomics 3' V2). Color denotes cell populations annotated by SingleR with malignant cells detected by CopyKat. (**B**) Distribution of pathway scores in the main cell populations. (left to right) cell populations were sorted by median pathway scores higher to lower. Eight out of 11 pathways are shown consisting of five to 22 unique downstream target molecules per pathway (MYC, MITF, EGFR, SP1, ERBB2, SERBP1, SOX2, CCND1). Meta-pathway represents a conglomerate expression score of downstream target molecules from all pathways combined. (**C**) Comparison of tumor cell-expressing pathway scores between low and high T cell-infiltrated tumors. Each data point represents one malignant cell. *n*=4249 and 495 malignant cells shown from low and high T cell-infiltrated tumors, respectively. (**D**) Heatmap showing the decreased tumor cell expression of the 108 genes enriched in IFN stimulating functions (same genes as shown in **Fig. 3D**; **Table S7**) in low vs high-T cell-infiltrated AM samples. P-values were computed by two-sided Two-Sample *t*-test in **C**, with denotation: * P <0.05; ** P <0.01; *** P <0.001 after FDR adjustment for multiple comparisons. NK = neutral killer cells.

#### Supplementary Tables

The **Supplementary Tables 1 to 7** were supplied as separate spreadsheets. Titles are listed below.

**Supplementary Table 1. Demographic and clinical characteristics of the patients diagnosed with AM in UPMC cohort.**

**Supplementary Table 2. List of whole transcriptomics, single-cell, and ICI-treated cohorts included in this study.**

**Supplementary Table 3. List of DEGs shared among UPMC, NW, and MIA AM cohorts comparing non-T cell-inflamed versus T cell-inflamed tumors**. “up” or “down” denotes the directionality of gene expression change in non-T cell-inflamed relative to T cell-inflamed tumors within each cohort. “unchanged’ indicates the DEGs did not pass filters of at least 1.5-fold change or had p-values larger than 0.05.

**Supplementary Table 4. Upstream regulator (pathways) predicted to be activated in non-T cell-inflamed tumors relative to T cell-inflamed tumors from the UPMC cohort.**

**Supplementary Table 5. Upstream regulator (pathways) predicted to be activated in non-T cell-inflamed tumors relative to T cell-inflamed tumors from the NW cohort.**

**Supplementary Table 6. Upstream regulator (pathways) predicted to be activated in non-T cell-inflamed tumors relative to T cell-inflamed tumors from the MIA cohort.**

**Supplementary Table 7. List of 108 genes enriched in IFN stimulating functions with decreased gene expression in low vs high-T cell-infiltrated AM tumors based on single-cell RNAseq data.**
